# Spatial, temporal and taxonomic patterns of insect extinction in Germany

**DOI:** 10.1101/2022.12.19.521006

**Authors:** Caspar A. Hallmann, Thomas Hörren, Axel Ssymank, Hubert Sumser, Heinz Schwan, Werner Stenmans, Mareike Vischer-Leopold, Livia Schäffler, Martin Sorg

## Abstract

Red lists represent an important instrument for evaluating the decline of species in space and time, for improving decisionmaking and for guiding conservation planning. However, globally, only a fraction of species has been categorized according to a red list, even in countries where insects are relatively well-studied. Such large knowledge gaps hinder conservation planning and ultimately jeopardize the maintenance of ecosystem functions. Given the recent reports on severe insect decline, it is now more than ever of great importance to obtain a reliable complete picture of the state of insects. We here derive an estimate of extinction rates and of the proportion of threatened species for the total insect community in Germany, and asses spatial and temporal of extinction patterns.

We found a regional extinction rate of 4.5% (1773-1937 species) for the area of Germany. Among extant insect species, 6% are classified as critically endangered (1856-2024 species), while among remaining species, a staggering 36.1% (10758-11086 species) is classified as threatened.

Higher trophic levels of zoophagous insects are often more sensitive to negative environmental changes due to their position in the food web, and at the same time are underrepresented in Red Lists. They are therefore disproportionately affected by these knowledge gaps.

This concerns particularly parasitoids which are taxa of regulatory importance and often higher extinction risk levels due to their trophic position.

Exemplary examination of the spatial scaling of red list categories indicate a far higher rate and risk and exemplary over ten times higher regional extinction rate when the reference area is gradually scaled down.

This illustrates the actual situation regarding the magnitude of regional species extinction events and extirpation risks that we have to assume for certain parts of the reference areas.

For a given region, the loss of the gene pool of populations specially adapted to a given region usually represents an irreversible biodiversity loss. In order to avoid further irreparable damage, the species threatened with extinction must be preserved with top priority. There is thus a considerable need for research in order to assess the conservation status of more than 56% of the insect species diversity in Germany and to immediately achieve a more balanced trait group representation in red lists.

## Introduction

Red lists represent an important instrument for evaluating the decline of species in space and time, for improving decisionmaking and for guiding conservation planning. However, globally, only a fraction of insect-species have been categorized according to a red list, even in countries where insects are well-studied such as in Germany. This large knowledge gap hinders conservation planning and ultimately jeopardizes effective conservation of species and ecosystem functioning. Given the recent reports on severe insect declines (1–6), obtaining a reliable and complete picture of the conservation status of insects, has become highly urgent.

Insects are an extremely important indicator group for the vast majority of ecosystems, both in aquatic and terrestrial habitats. Owing to their great diversity, they naturally exhibit a myriad of ecological functions. However, many of the ecological functions attached to species characteristics are not well understood. Coupled with an insufficient knowledge on conservation status of species, this lack of understanding further restricts a comprehensive assessment of ecosystem functions at risk, and prohibits targeted conservation strategies.

Furthermore, while arguably the status of several, usually charismatic taxa among insects may be covered at national levels and beyond, knowledge on their status at smaller spatial scales is often even less comprehensive - even though that is the typical spatial scale of management actions. As such, we are in need of a cross-scale analysis of the conservation status of species, in order to identify local conservation priorities, and to provide management recommendations.

We assemble and summarize current knowledge on conservation status of insects in Germany, in order to detect and describe the aforementioned knowledge gaps. Our objectives are x-fold. First, we account for the incompleteness of red list categorization, and provide estimates of the expected (complete) number of regionally extinct, endangered and threatened species, while accounting for family-level species traits (7). Second, we compiled red list data sources across different spatial scales from local district to continent, and examined the scale dependent distribution of extinction rate and number of extinct species. Third, we examine temporal patterns in extinction and cumulative extinction rates at national level. Our work serves to gain an overall overview of the current state of knowledge regarding the conservation status of insects, and to identify knowledge gaps as well as priorities for required actions.

## Methods

### red list data sources

We consider the sources mentioned in Supplement 1 for national and regional red lists for Germany, North Rhine Westphalia and the lower rhine region. For the smallest study area in the region around the city of Krefeld, were compiled from long term presence-absence records as collected by the Entomological Society of Krefeld.

### Species diversity and red lists of Germany

Past accounts (8, 9) have resulted in estimates of number of insects present in Germany to exceed 33,000. Thus, this taxonomic class comprises at least 7-8 times more species that all vertebrates and plants together. The German Barcode of Life (GBOL) reference library curretnly accounts for at least one thousand more insect species (34,085), and is thus regarded a more comprehensive assessment. However, the red list of insects is by far not as comprehensive as those of vertebrates and plants, with include 14,940 species in total (43.8% compared to total species numbers of GBOL). Moreover, the red listing includes only ten out of 25 insect orders known to be present in Germany (Fig. 1), while within orders, species coverage ranges between 0% for species rich insect families and up to 100% for species-poor groups (Fig. 1).

**Fig. 1.**
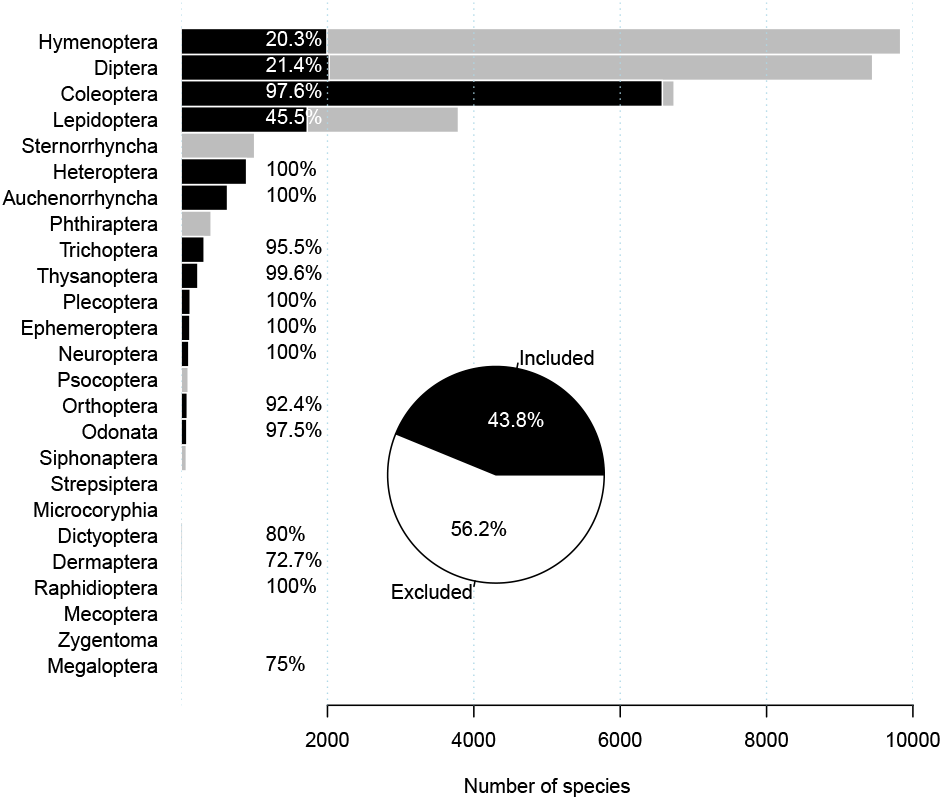
Number of insect species per order known to exists in Germany, as well as number of species categorized in the red list of Germany.

Out of the ten insect orders included in the red list assessment of Germany, there may still be species that do not have an appropriate conservation status because for example they have been classified as data-deficient (Fig. 2). For those species for which a formal red list assessment is given, on average, 4.1% are classified as extinct, 5.7% are classified as threatened with extinction, 8.7% as highly endangered, 10.2% as endangered and 45.8% are classified as non-threatened.

**Fig. 2.**
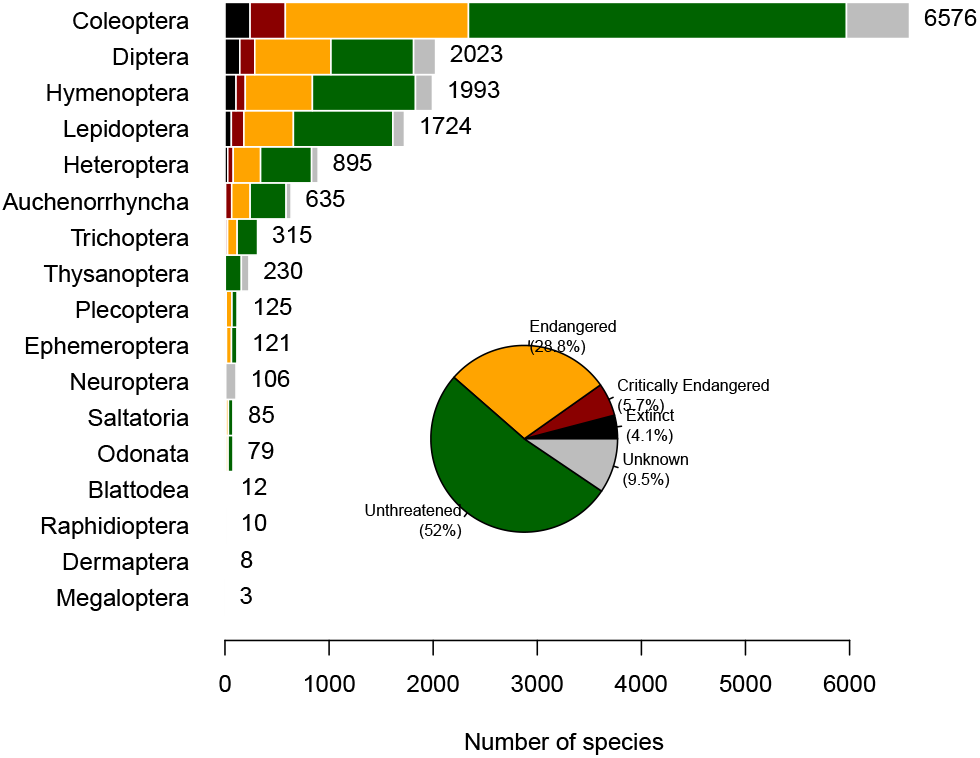
Simplified distribution of red list categories among ten insect orders in Germany. Critically endangered consists of category 1, Endangered are the combined classes of threat categories 2, 3, R and G, not threatened are the categories V and *, and unknown include the categories data deficient or unclassified species.

### Statistical analysis

First we provide estimates of the numbers of extinct, endangered and threatened species, while accounting for incompleteness of the national red list assesment, as well as trophic and habitat characteristics of the species (at family level). Second, we derive empirical estimates of scale dependent extinction, and scale depended number of extinct species. Third, we derive estimates of temporal extinction at national level.

#### Models of risk categories

We start by stating that the number of species included in each red list category follow a multinomial distribution, with *K* number of species categorized (presently 14,940), and cell probabilities 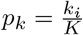, where *k_i_* the number of species in each category *i*. For practical reasons however, it suffices to reduce the problem to two independent binomial models: that of extinction, and that of the threatened versus non-threatened for extant species. The goal is derive the number of species (*m*) in each category (extinct or threatened) for the full number of species in Germany (*M*), with corresponding unknown probabilities *p_m_*.

We derive four models to impute red list categories for nonclassified insect species. Our first model (the *null-model)* assumes no variation in categorization among species, or other grouping (e.g. taxonomic or functional), in the rate of categories, i.e. *p_k_* = *p_m_*. In other words, our null-model assumes that the insect species included in the red list assessment are a representative sample of all insect species present in Germany. The assumption implies that the included species within a particular insect order are representative to all species within that particular order, and that included orders are representative to non-included orders. Our model reads

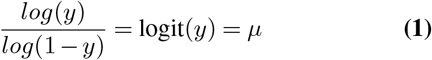

In a two subsequent models, we attempt to relax to some degree the assumption of equal *p_k_* probabilities among insect orders by including potential explanatory variables at order of the family level. We allow fraction of species in each category to be species-richness dependent. Here, the assumption is made that the number of species in each order has a deterministic relation to the fraction of species that have gone extinct, or are threatened, besides empirical justification, is presently also motivated by examination of the data at hand (Figure 2). Our second model reads

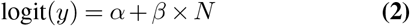

where *N* the vector with number of species in each family of the corresponding species.

In a third model, we examine whether number of species in each red list category, and corresponding probabilities, are related to family-level traits, such as diet and habitat preferences. To this end we use a recently compiled database of these traits for all insect families in Germany (7), allowing us to both model extinction and threatened status probabilities for red listed species, as well as to predict status probabilities for non-red listed species. This newly assembled trait-database is categorized using a fuzzy-classification at family level, in essence providing a level of association between a given family with the various trait categories.

We run a number of models. Our full model is expressed as

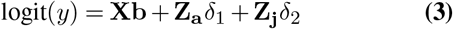

where y the binary response of extinction per species, **X** the row-proportionalized matrix (i.e. ∑ *X* [*i*,.] = 1) of larval food preference, **b** a vector of corresponding coefficients for the log-odds of association to trait categories with respect to larval food preference, while *Z_a_* and *Z_b_* indicator variables of aquatic life history in adult and larval life stages respectively. In all three models we rely on binomial distributions, with a *logit*-link embeded in a generalized linear model. In a final step, models were compared by Akaikes Information Criterion (AIC) in order to identify the most parsimonious representation.

#### Spatial scale of categorization

It is widely known that extinction is scale dependent, with regional extinction processes decreasing with larger reference area of assessment. To examine this relationship, we compiled the available data on insect extinctions at different spatial scales, from continent (Europe), to local (municipality), and derived empirical scale dependent estimates of extinction. Next, we derived species accumulation curves (SAR), and integrated them with the scale dependent extinction probabilities, to produce scale dependent estimates of number of extinct species.

1. Species area curves (SAR) are usually characterized by a curvi-linear relationship: *S* = *cA^z^*, where *S* is the number of species, *A* the area and *c* and *z* constants
2. Extinction risk of a species is inversely related to population size. That is, for given mean extirpation rate (*r*) per local population, the extinction probability (over a larger geographical area) decreases with increasing area (and number of populations).
3. Population size is linearly dependent on area.

#### Temporal rate of regional extinction

Red list data for extinct species include the last year of a recording. Under the assumption that the particular year the species is last seen is the year of extirpation, we examined temporal patterns in extinction risk. In particular, we were interested in non-linearity patterns over time.

## Results

### Distribution of traits

The species distribution of trophic trait categories included in the red list assessment differed considerably compared to the composition of the total list of species known to be present in Germany. In particular, phytophagous species are generally over represented in the red list assessment, contrary to zoophagous and detritophagous species that are under represented (Fig. 3A). Species in aquatic environments are overrepresented in the red list as compared to the national species list (Fig. 3B). Within zoophagous species, red list assessments of species are strongly biased towards predatory species, with a clear under-representation of parasitoid species (Fig. 3C), while within phytophagous species, red-list representation is biased towards phyllophagous species, with a complete absence of gall-inducing species and miners (Fig. 3D).

**Fig. 3.**
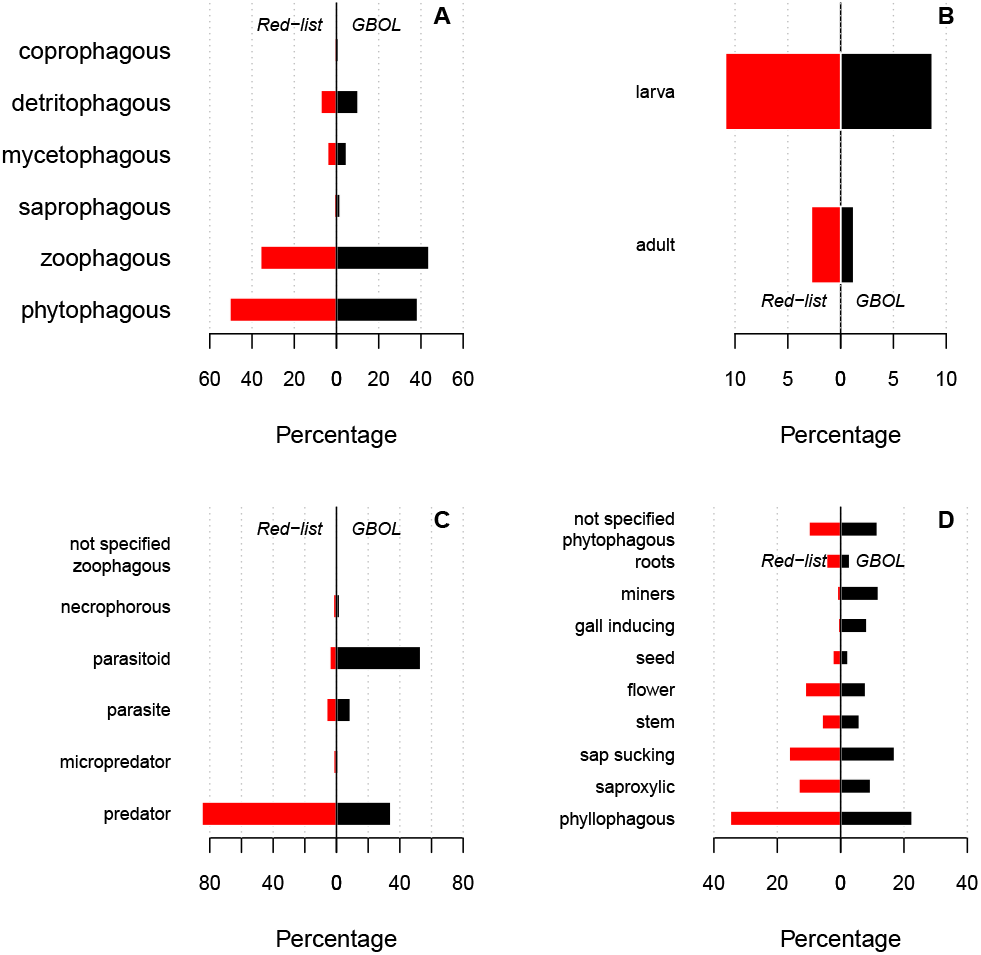
Comparison of trait distribution representation between red listed species and total known species (GBOL database). A: Relative percentage of larval feeding strategy. B: Percentage species in terrestrial environment for larvae. C: Percentage feeding specialization for zoophagous species. D: Percentage feeding specialization for phytophagous species.

### Models for red list extinction, endangerment and threatened status

Among ten competing models, models including subcategories of larval-diet were more supported by the data as compared to models including major diet categories only. This was found to be true across the three response variables i.e. fraction extinct, fraction critically endangered, and fraction threatened (Supplementary tables S1–S3). In particular, most parsimonious models were found to be those including subcategories larval diet class and an indicator of association to aquatic vs terrestrial realm for extinction and threatened status categories.

The largest fraction of extinct species was found to be among coprophagous species (Fig. 4), followed by zoophagous species. The highest fraction critically endangered species to be present among phyllophagous species specializing on roots, although on average a higher fraction of coprophagous species was classified as critically endangered, as compared to phyllophagous species (Fig. 5). Estimates of model coefficients of the most parsimonious model for each response variable are given in Supplementary tables S4–S6.

**Fig. 4.**
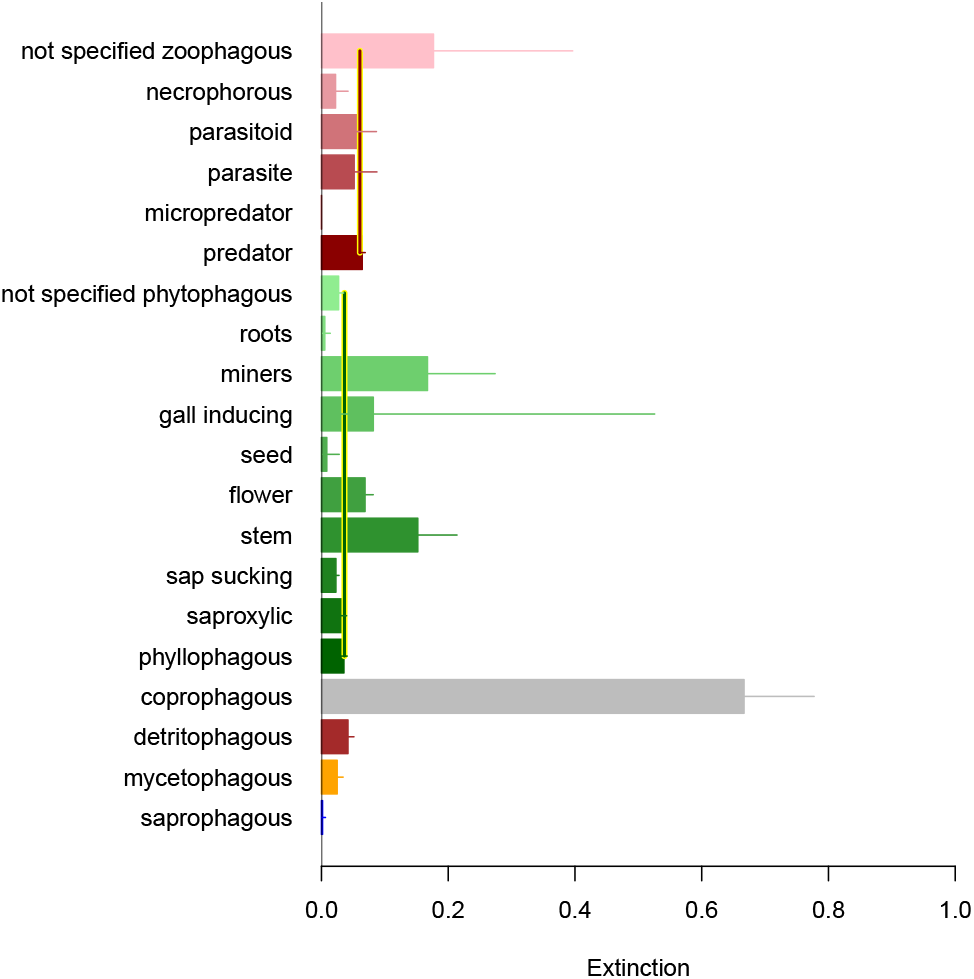
Estimated fraction of extinct species across different larval feeding strategies. Vertical lines represent the averages across phyllophagous and zoophagous species.

**Fig. 5.**
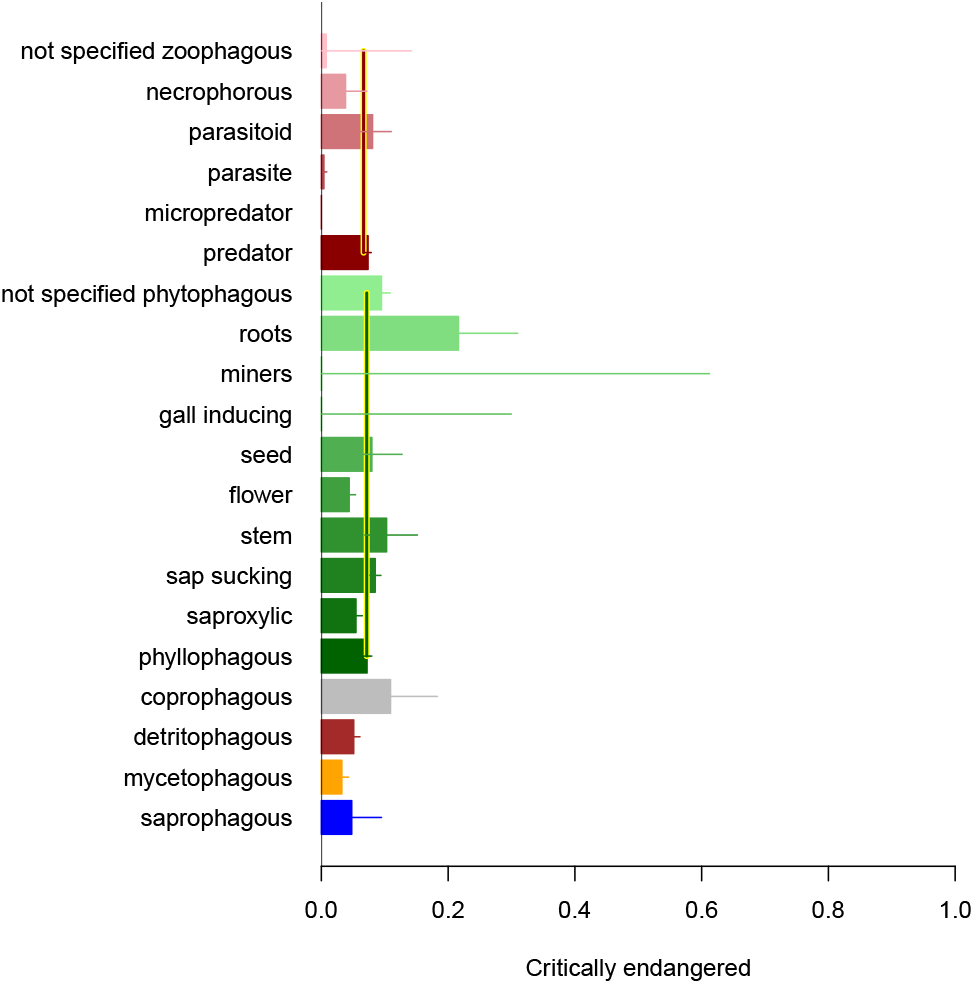
Estimated fraction of critically endangered species across different larval feeding strategies. Vertical lines represent the averages across phyllophagous and zoophagous species.

### Models of scale dependence

The species-area curve fitted to the present insect data showed results consistent with global results, with best estimates *y* = *α* × *A^z^* = *exp*(1.7862 + *log*(*Area*) × 0.2527). The exponent coefficient *z* was found to be well within the reported global *z*–values among other life forms (i.e. 0.2-0.3) (). Among insect species groups, the random intercept effect coefficient was found to be *ϵ* = 1.369 (on the log scale).

We found a strong scale dependent extinction probability across 30 insect (Fig. 6), with a significantly negative coefficient for log-area (*logit*(*y*) = 2.3180 – 0.4598 × *log*(*Area*), *t*-value=-21.572, p-value < 0.001). Between species random effect variation was estimated at *ϵ* = 0.7361.

**Fig. 6.**
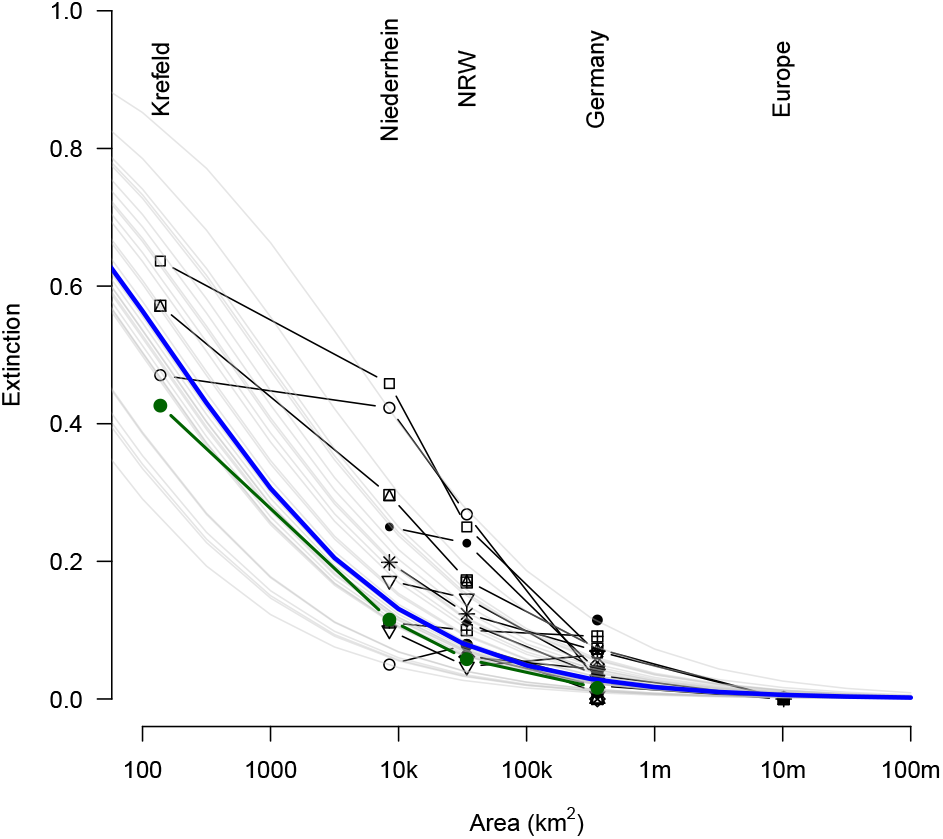
Estimated mean scale-dependent extinction probability for insects. Points and black lines represent individual insect groups. Grey lines are fitted group-dependent probabilities while blue line represent mean probability across all insect groups. The green line represent scale dependent plant species extinction probabilities.

Integrating scale dependent extinction probability with the species area curve, allowed us to examine scale dependent number of extinct species. We found a humped shape relationship of number of species that are extinct as function of scale (Fig. 6), suggesting that most species have gone extinct at above-minimum of the area range considered presently, around 300 km^2^.

#### Temporal rate of regional extinction

We found a highly nonlinear extinction rate over time (1800-1970), with low extinction rates in the period 1800-1940, followed by much higher extinction rates in the following period.

**Table 1.** Table with estimates of number of extinct, endangered and threatened species, given model formulation, as well as 95% confidence intervals of binomial variation. Values in bold are those of the most parsimonious models.

| model | Extinct | Endangered | Threatened |
| --- | --- | --- | --- |
| $\mu$ | 1535(1460-1610) | 2131(2044-2219) | 10746(10583-10909) |
| N | 1553(1478-1629) | 1977(1893-2062) | 10225(10064-10386) |
| Feeding | 1614(1538-1691) | 2074(1988-2161) | 10758(10595-10921) |
| Feeding + $A_{qad}$ | 1644(1567-1722) | 2062(1976-2148) | 10801(10638-10965) |
| Feeding + $A_{qla}$ | 1647(1570-1725) | 2048(1963-2135) | 10730(10567-10893) |
| Feeding + $A_{qad}$ + $A_{qla}$ | 1645(1567-1723) | 2049(1963-2135) | 10727(10564-10890) |
| Extended feeding | 1858(1777-1941) | **1940(1856-2024)** | 10795(10632-10958) |
| Extended feeding + $A_{qad}$ | **1855(1773-1937)** | 1939(1856-2023) | **10803(10640-10966)** |
| Extended feeding + $A_{qla}$ | 1859(1777-1941) | 1938(1854-2022) | 10803(10640-10967) |
| Extended feeding + $A_{qad}$ + $A_{qla}$ | 1854(1772-1936) | 1939(1856-2023) | 10782(10619-10946) |

## Discussion

We found that Red-list categorization is unbalanced with respect to the taxonomic and functional insect diversity in Germany (Fig. 1). In addition, we found that extinction rates, as well as the fraction of critically endangered species, strongly dependent on the trophic position of the species. In turn, extinction and endangerment levels cannot be simply extrapolated to the national species list, based on the red list proportions alone. Among the about 34,000 species known to be or have been present in Germany, and taking into account taxonomic and functional uncertainty, we estimate 1855 species to have gone extinct, and 1940 species to be threatened with extinction, which is the highest endangerment category of the Red List in Germany.

The by far biggest, identified gaps in knowledge with respect to conservation-status assessments, are to be found among the very species-rich parasitoid Hymenoptera, and a larger number of the Diptera families (Fig. 1). For these two orders alone, a remarkable 14,500 species do not presently have a conservation assessment in place.

Knowledge on the status of insects in Germany is therefore far from complete.

For a better understanding of regional and national biodiversity changes in the time line, a comprehensive threat assessment of as many species as possible is needed. Priority in data collection and vulnerability assessment should be given to taxa with the most identified knowledge deficits.

In insects this is especially necessary for species-rich families with parasitoid lifestyles.

We found a strong sclae dependent extinction probability of insect species, resulting in a humped pattern of number of extinct species with increasing area. This humped relationship may theoretically arise as result of contracting species ranges, suggesting habitat loss and fragmentation hindering recolonization.

We found that extinction rates were accelerating over time, with higher total extinction numbers coinciding with the industrialization period after the second world war.

By definition, the Red Lists cannot contribute to the detection of current extinction processes (i.e of recent decades), since a species is considered extinct only in the absence of any record over time period of several decades Hence, present results based on the red-list data at hand, are unable to provide an up to date assessment of current extinction rates within the last decades. However, recently documented insect declines in Germany (1–3) render a decline in extinction rates in recent years highly unlikely. Our estimated extinction rates and numbers of extinct species should therefore be considered a conservative assessment.

## Recommendations

Our results refer to a country with relatively high land use intensity, population density and industrialization. On the other hand, there is a comparatively high level of knowledge about insect diversity and nature protection in Germany, based on a long tradition of entomological research, nature conservation policy and research-funding potential.

Owing to the data red lists are build upon, this is a view of the past and not a reflection about events of the last two decades. Our results show the best possible current approximation to the reality of biodiversity loss that has occurred in the reference area and exemplary sub-areas.

We characterize regional extinction events as irreversible damage if they result in the loss of population characteristics anchored in their regional gene pool.

The conservation of stable (meta)populations of species threatened with extinction therefore as a top priority goal.

Insect species in the category threatened with extinction (Category 1 of the red list of Germany) usually have a requirement profile that is no longer met in the “normal landscape” in Germany. As a rule, the last populations of these species are located in nature reserves and especially in the Natura 2000 network of protected areas of the European Union. These species are also often identified as indicators of a higher quality condition of strictly protected habitats of the EU Flora Fauna Habitat Directive.

In order to be able to comprehensively assess biodiversity change and to provide meaningful conservation and management recommendations, we need to be informed about exactly those species. As described earlier, this is is currently not the case at the national level, and even less so at smaller spatial scales.

At least for further exemplary sub-regions, it is therefore necessary to close these knowledge gaps in order to better understand what actually happens in which dimensions when the area is scaled down. This knowledge should be available for the overall diversity of insects within a network of protected areas selected as representative examples. To this end, thorough investigations should be initiated in order to responsibly build up sufficient knowledge and take targeted protective measures.

## Conclusions

### Future directions

Closing the knowledge gaps for these higher taxa was hampered in the past by the small or regionally non-existent number of entomologists who are able to identify species of these insect families by conventional taxonomic methodology.

The decline in the total number of flying insects was determined using the methodology of standardized Malaise trapping, which offer opportunity to collect sufficient data for particularly these insect orders (Hymenoptera, Diptera).

The hope of closing existing knowledge gaps additionally lies in the application of genetic methods of species determination (metabarcoding) to the mixed samples of such efficient detection methods. Current knowledge suggests that standardized malaise traps, in combination with genetic methods for species identification, are particularly well suited to fill most of these identified gaps.

## ACKNOWLEDGEMENTS

Funded by the German Federal Ministry for the Environment, Nature Conservation and Nuclear Safety (BMUV), handled by the The German Federal Agency for Nature Conservation (BfN), grant number FKZ 3516850400. We would also like to thank many experts from the Entomological Society Krefeld for advice and assistance during the processing of the expert assessments.

## Author contributions

TH, CH and MS developed the conceptual framework. TH and MS collected and compiled the data. CH, TH and MS drafted the first version of the manuscript. All authors contributed to the article, approved the version to be published and agreed to be accountable for all aspects of the work.

## Supplement 1

References of red list publications used .pdf file: http://entomologica.org/dl/rl-publ.pdf

## Supplement 2

Compiled red lists for the insect fauna of Germany .pdf file: http://entomologica.org/dl/rl-germany.pdf

## Supplement 3

Table for comparing the red lists of Germany with the area of North Rhine-Westphalia, the lower Rhineland and the area around Krefeld for selected Taxa .pdf file: http://entomologica.org/dl/rl-s3.pdf

## Supplement 4 - Supplementary tables

**Table S1.** Table of results of extinction risk for various model formulations. *μ* is a simple mean, N is number of species in family, Feeding is the larval feeding strategy (See Fig 3A) and *Aq_ad_* and *Aq_la_* represent effects of adult and larval association to aquatic environment respectively. Extended feeding is the larval feeding strategy whereby the phytophagous and zoophagous strategies are further specified in subclasses (See Figs. 3D and 3E).

| model | df | AIC | deviance | $\Delta$ AIC | weight |
| --- | --- | --- | --- | --- | --- |
| $\mu$ | 1.00 | 4968.63 | 4966.63 | 90.45 | 0.00 |
| N | 2.00 | 4970.49 | 4966.49 | 92.31 | 0.00 |
| Feeding | 6.00 | 4913.77 | 4901.77 | 35.59 | 0.00 |
| Feeding + $A_{qad}$ | 7.00 | 4905.92 | 4891.92 | 27.74 | 0.00 |
| Feeding + $A_{qla}$ | 7.00 | 4912.85 | 4898.85 | 34.67 | 0.00 |
| Feeding + $A_{qad}$ + $A_{qla}$ | 8.00 | 4907.92 | 4891.92 | 29.74 | 0.00 |
| Extended feeding | 20.00 | 4886.24 | 4846.24 | 8.06 | 0.01 |
| **Extended feeding + $A_{qad}$** | **21.00** | **4878.18** | **4836.18** | **0.00** | **0.69** |
| Extended feeding + $A_{qla}$ | 21.00 | 4886.59 | 4844.59 | 8.41 | 0.01 |
| Extended feeding + $A_{qad}$ + $A_{qla}$ | 22.00 | 4879.92 | 4835.92 | 1.74 | 0.29 |

**Table S2.** Table of results of endangerement risk for various model formulations. *μ* is a simple mean, N is number of species in family, Feeding is the larval feeding strategy (See Fig 3A) and *Aq_ad_* and *Aq_la_* represent effects of adult and larval association to aquatic environment respectively. Extended feeding i sthe larval feeding strategy whereby the phytophagous and zoophagous strategies are further specified in subclasses (See Figs. 3D and 3E).

| model | df | AIC | deviance | $\Delta$ AIC | weight |
| --- | --- | --- | --- | --- | --- |
| $\mu$ | 1.00 | 6291.81 | 6289.81 | 73.97 | 0.00 |
| N | 2.00 | 6276.89 | 6272.89 | 59.05 | 0.00 |
| Feeding | 6.00 | 6279.47 | 6267.47 | 61.63 | 0.00 |
| Feeding + $A_{qad}$ | 7.00 | 6280.17 | 6266.17 | 62.33 | 0.00 |
| Feeding + $A_{qla}$ | 7.00 | 6279.67 | 6265.67 | 61.83 | 0.00 |
| Feeding + $A_{qad}$ + $A_{qla}$ | 8.00 | 6281.47 | 6265.47 | 63.64 | 0.00 |
| **Extended feeding** | **20.00** | **6217.84** | **6177.84** | **0.00** | **0.51** |
| Extended feeding + $A_{qad}$ | 21.00 | 6219.55 | 6177.55 | 1.71 | 0.22 |
| Extended feeding + $A_{qla}$ | 21.00 | 6219.73 | 6177.73 | 1.89 | 0.20 |
| Extended feeding + $A_{qad}$ + $A_{qla}$ | 22.00 | 6221.55 | 6177.55 | 3.71 | 0.08 |

**Table S3.** Table of results of threatened risk for various model formulations. *μ* is a simple mean, N is number of species in family, Feeding is the larval feeding strategy (See Fig 3A) and *Aq_ad_* and *Aq_la_* represent effects of adult and larval association to aquatic environment respectively. Extended feeding i sthe larval feeding strategy whereby the phytophagous and zoophagous strategies are further specified in subclasses (See Figs. 3D and 3E).

| model | df | AIC | deviance | $\Delta$ AIC | weight |
| --- | --- | --- | --- | --- | --- |
| $\mu$ | 1.00 | 15715.42 | 15713.42 | 170.59 | 0.00 |
| N | 2.00 | 15676.53 | 15672.53 | 131.70 | 0.00 |
| Feeding | 6.00 | 15671.79 | 15659.79 | 126.96 | 0.00 |
| Feeding + $A_{qad}$ | 7.00 | 15669.59 | 15655.59 | 124.76 | 0.00 |
| Feeding + $A_{qla}$ | 7.00 | 15673.28 | 15659.28 | 128.46 | 0.00 |
| Feeding + $A_{qad}$ + $A_{qla}$ | 8.00 | 15666.66 | 15650.66 | 121.84 | 0.00 |
| Extended feeding | 20.00 | 15551.12 | 15511.12 | 6.29 | 0.02 |
| **Extended feeding + $A_{qad}$** | **21.00** | **15544.83** | **15502.83** | **0.00** | **0.54** |
| Extended feeding + $A_{qla}$ | 21.00 | 15552.80 | 15510.80 | 7.98 | 0.01 |
| Extended feeding + $A_{qad}$ + $A_{qla}$ | 22.00 | 15545.26 | 15501.26 | 0.44 | 0.43 |

**Table S4.** Tabel of GLM-model coefficients for most parsimonious model for extinction risk

|  | Estimate | Std. Error | z value | Pr(>\|z\|) |
| --- | --- | --- | --- | --- |
| saprophagous | -6.328 | 1.270 | -4.981 | 0.000 |
| mycetophagous | -3.671 | 0.329 | -11.143 | 0.000 |
| detritophagous | -3.131 | 0.213 | -14.725 | 0.000 |
| coprophagous | 0.694 | 0.557 | 1.246 | 0.213 |
| phyllophagous | -3.303 | 0.137 | -24.074 | 0.000 |
| saproxyllic | -3.412 | 0.223 | -15.302 | 0.000 |
| sap sucking | -3.759 | 0.199 | -18.857 | 0.000 |
| stem | -1.719 | 0.417 | -4.120 | 0.000 |
| flower | -2.606 | 0.186 | -14.032 | 0.000 |
| seed | -4.766 | 1.218 | -3.912 | 0.000 |
| gall inducing | -2.418 | 2.521 | -0.959 | 0.337 |
| miners | -1.605 | 0.630 | -2.546 | 0.011 |
| roots | -5.246 | 0.982 | -5.342 | 0.000 |
| not specified phytophagous | -3.581 | 0.263 | -13.597 | 0.000 |
| predator | -2.677 | 0.075 | -35.574 | 0.000 |
| micropredator | -27.285 | 467.607 | -0.058 | 0.953 |
| parasite | -2.911 | 0.567 | -5.137 | 0.000 |
| parasitoid | -2.734 | 0.379 | -7.206 | 0.000 |
| necrophorous | -3.782 | 0.654 | -5.784 | 0.000 |
| not specified zoophagous | -1.538 | 1.118 | -1.376 | 0.169 |
| adult_aquatic | -1.005 | 0.366 | -2.745 | 0.006 |

**Table S5.** Tabel of GLM-model coefficients for most parsimonious model for endangerment risk

|  | Estimate | Std. Error | z value | Pr(>\|z\|) |
| --- | --- | --- | --- | --- |
| saprophagous | -3.000 | 0.735 | -4.080 | 0.000 |
| mycetophagous | -3.401 | 0.296 | -11.491 | 0.000 |
| detritophagous | -2.902 | 0.182 | -15.987 | 0.000 |
| coprophagous | -2.104 | 0.603 | -3.486 | 0.000 |
| phyllophagous | -2.555 | 0.105 | -24.391 | 0.000 |
| saproxylis | -2.854 | 0.180 | -15.856 | 0.000 |
| sap sucking | -2.377 | 0.111 | -21.512 | 0.000 |
| stem | -2.171 | 0.443 | -4.904 | 0.000 |
| flower | -3.082 | 0.218 | -14.145 | 0.000 |
| seed | -2.446 | 0.523 | -4.676 | 0.000 |
| gall inducing | -10.942 | 10.092 | -1.084 | 0.278 |
| miners | -9.007 | 9.463 | -0.952 | 0.341 |
| roots | -1.290 | 0.486 | -2.654 | 0.008 |
| not specified phytophagous | -2.254 | 0.150 | -15.010 | 0.000 |
| predator | -2.523 | 0.070 | -36.152 | 0.000 |
| micropredator | -24.983 | 283.790 | -0.088 | 0.930 |
| parasite | -5.544 | 0.753 | -7.362 | 0.000 |
| parasitoid | -2.436 | 0.345 | -7.066 | 0.000 |
| necrophorous | -3.221 | 0.667 | -4.830 | 0.000 |
| not specified zoophagous | -4.888 | 3.076 | -1.589 | 0.112 |

**Table S6.**
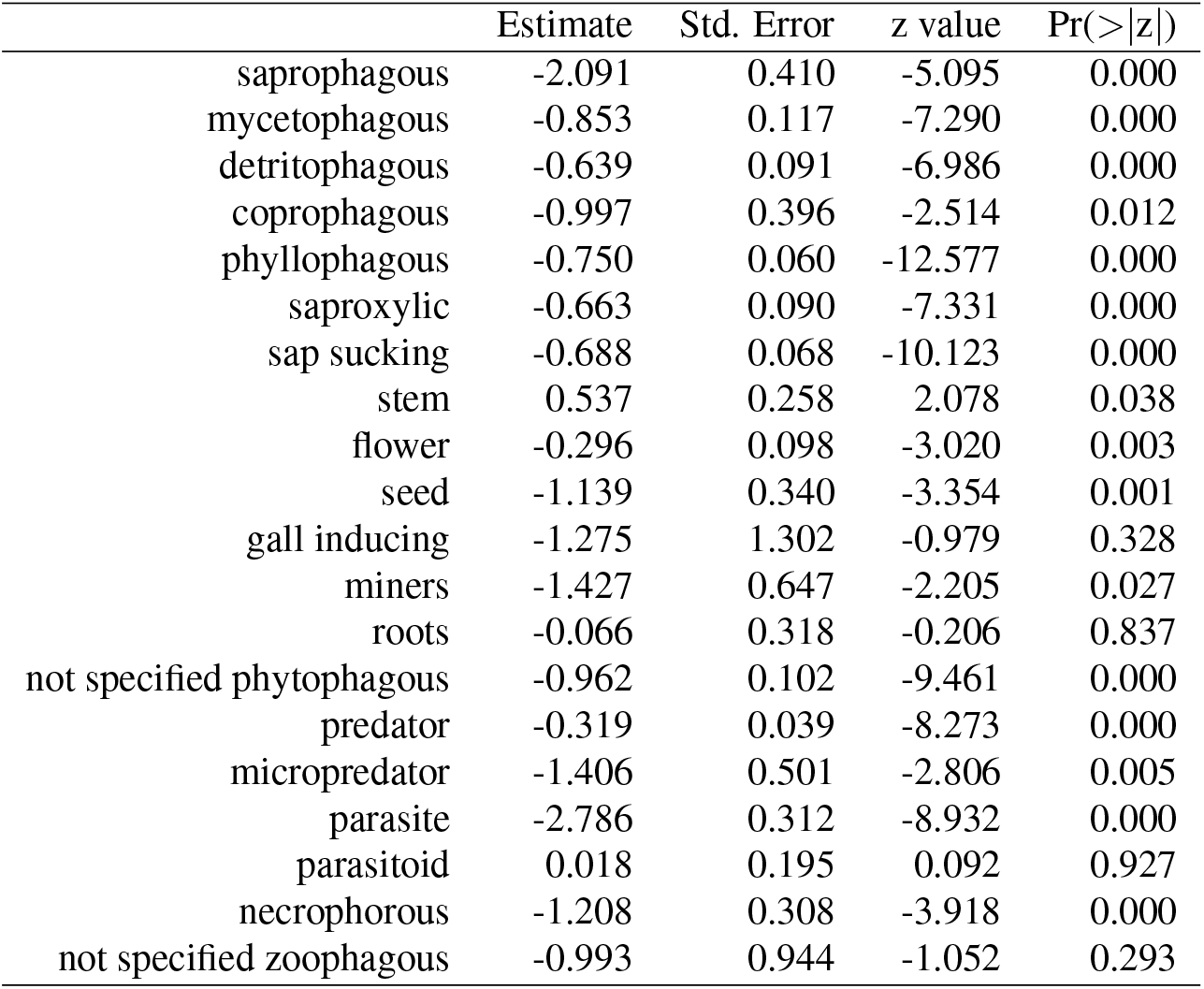
Tabel of GLM-model coefficients for most parsimonious model for threatened risk

## References

1. Caspar A Hallmann, Martin Sorg, Eelke Jongejans, Henk Siepel, Nick Hofland, Heinz Schwan, Werner Stenmans, Andreas Müller, Hubert Sumser, Thomas Hörren, et al. More than 75 percent decline over 27 years in total flying insect biomass in protected areas. PloS one, 12(10):e0185809, 2017.

2. Sebastian Seibold, Martin M Gossner, Nadja K Simons, Nico Blüthgen, Jörg Müller, Didem Ambarli, Christian Ammer, Jürgen Bauhus, Markus Fischer, Jan C Habel, et al. Arthropod decline in grasslands and forests is associated with landscape-level drivers. Nature, 574 (7780):671–674, 2019.

3. Caspar A Hallmann, Axel Ssymank, Martin Sorg, Hans de Kroon, and Eelke Jongejans. Insect biomass decline scaled to species diversity: General patterns derived from a hoverfly community. Proceedings of the National Academy of Sciences, 118(2), 2021.

4. Roel Van Klink, Diana E Bowler, Konstantin B Gongalsky, Ann B Swengel, Alessandro Gentile, and Jonathan M Chase. Meta-analysis reveals declines in terrestrial but increases in freshwater insect abundances. Science, 368(6489):417–420, 2020.

5. David L Wagner, Eliza M Grames, Matthew L Forister, May R Berenbaum, and David Stopak. Insect decline in the anthropocene: Death by a thousand cuts. Proceedings of the National Academy of Sciences, 118(2), 2021.

6. Pedro Cardoso, Philip S Barton, Klaus Birkhofer, Filipe Chichorro, Charl Deacon, Thomas Fartmann, Caroline S Fukushima, René Gaigher, Jan C Habel, Caspar A Hallmann, et al. Scientists’ warning to humanity on insect extinctions. Biological Conservation, 242:108426, 2020.

7. Thomas Hörren, Martin Sorg, C.A Hallmann, Axel Ssymank, Niklas W. Noll, Livia Schäffler, and Christoph Scherber. A universal insect trait tool (itt, v.1.0) for statistical analysis and evaluation of biodiversity research data. BioRxiv, 2022.

8. Wolfgang Völkl, Theo Blick, Paul M Kornacker, and Harald Martens. Quantitativer überblick über die rezente fauna von deutschland. Mollusca, 1:4, 2004.

9. B Klausnitzer. Die insektenfauna deutschlands (entomofauna germanica) - ein gesamtüberblick. Linzer Biologische Beitraege, 37(1):87—97, 2005.

